# Defining coarse-grainability in a model of structured microbial ecosystems

**DOI:** 10.1101/2021.07.17.452786

**Authors:** Jacob Moran, Mikhail Tikhonov

## Abstract

Despite their complexity, microbial ecosystems appear to be at least partially “coarse-grainable” in that some properties of interest can be adequately described by effective models of dimension much smaller than the number of interacting lineages (frequently in the dozens or hundreds). This is especially puzzling since recent studies demonstrate that a surprising amount of functionally relevant diversity is present at all levels of resolution, down to strains differing by 100 nucleotides or fewer. Rigorously defining coarse-grainability and understanding the conditions for its emergence is of critical importance for understanding microbial ecosystems. To begin addressing these questions, we propose a minimal model for investigating hierarchically structured ecosystems within the framework of resource competition. We use our model to operationally define coarse-graining quality based on reproducibility of the outcomes of a specified experiment and show that a coarse-graining can be operationally valid despite grouping together functionally diverse strains. We further show that, at least within our model, a high diversity of strains (while nominally more complex) may in fact facilitate coarse-grainability. However, this only applies if the pool of interacting strains is sampled from the appropriate “native” environment, as we discuss.

---

Microbial communities are complex dynamical systems composed of a highly diverse collection of interacting species, and yet they often appear to be at least partially “coarse-grainable”, meaning that some properties of interest can be predicted by effective models of dimension much smaller than the number of interacting lineages. For example, industrial bioreactors consisting of hundreds of species are well described by models with ≲ 10 functional classes [1, 2]. What makes this possible? One potential explanation is that coarse-grainability is a direct consequence of the hierarchically structured trait distribution across organisms. If 100 interacting phenotypes are all close variants of only 10 species, which can be further grouped into just two families, it is natural to expect that the diverse community might be approximately described by a 2- or 10-dimensional model. Under this view, effective models are possible because ecosystems are less diverse than a naïve counting of microscopic strains might suggest.

However, recent data reveal this intuition to be too simplistic: a surprising extent of relevant diversity persists at all levels of resolution. Numerous studies have highlighted the role of strain-level variation in shaping the functional repertoire of a microbial population [3–10]. A recent work by Goyal et al. concludes that strains might indeed be “the relevant unit of interaction and dynamics in microbiomes, not merely a descriptive detail” [11]. Surprisingly, however, a greater strain diversity can sometimes enhance predictability instead of undermining it [12]. Equally puzzling, the notion of a bacterial species is undoubtedly useful, despite collapsing together strains that famously may collectively share only 20% of their genes [13]. Moreover, by some assessments, the species-level characterization of a community appears to *be too detailed* and can be coarse-grained further, e.g., to the level of a taxonomic family [14].

Rigorously defining “coarse-grainability” and understanding the conditions for its emergence is of critical importance: harnessing coarse-grainability is our main instrument for understanding, predicting or controlling the behavior of these complex systems. Can an ecosystem be coarse-grainable for some purposes but not others? Or in some environments but not others? Can we ever expect the coarse-grained descriptions derived in the simplified environment of a laboratory to generalize to the complex natural conditions? Addressing this exciting set of general questions is an important challenge at the interface of theoretical microbial ecology and statistical physics.

Here, we introduce a theoretical framework to begin addressing these questions. The novelty of our approach is two-fold. First, we propose a minimal model for investigating hierarchically structured ecosystems. Much recent work studies the behavior of large microbial ecosystems in the unstructured regime, where the traits of interacting organisms are drawn randomly [e.g. 14–20]. However, real ecosystems assemble from pools of taxa whose trait distributions are highly non-random due to functional constraints, common selection pressures, or common descent. These factors create structure at all levels, from the distribution of genes across strains in microbial pangenomes [21–23] to the distribution of function across taxa [24, 25], with important implications for dynamics or responses to perturbations [26, 27]. In natural communities, taxa can often be grouped by identifiable functional roles, often represented by closely related species or strains. As we seek to define and characterize ecosystem coarse-grainability, it seems clear that this structure must play an important role. Our model implements such structure within a consumer-resource framework in a simple, principled way through trait interactions.

Our second novelty is a framework for defining and evaluating a hierarchy of coarse-grained descriptions. The ultimate performance criterion for a coarse-graining scheme would be its ability to serve as a basis for a predictive model, capable of predicting ecosystem dynamics or properties. However, finding the ‘most predictive model’ is a difficult problem. Here, as a simpler first step, we propose an operational approach which is inspired by the experiments of Ref. [14] and is based on the reproducibility of experimental outcomes. Specifically, we focus on a particular form of coarse-graining in which taxa are grouped together into putative functional groups. Grouping means omitting details, and we say that details are safe to ignore if they do not change the outcome of some specified experiment. Importantly, as we will show, choosing different experiments changes which, or whether, details can be ignored.

Specifically, we will define how ecosystems can be coarse-grainable in the weak sense, where a desired performance of a coarse-graining can be achieved in a given environment, and in the strong sense, where the performance of a given coarse-graining is *maintained* even as environment complexity is increased. We will demonstrate that the same ecosystem can be coarse-grainable under one criterion—even in the strong sense—and not at all coarse-grainable under another. This will reconcile the apparent paradox mentioned above, showing that a coarse-graining can be operationally valid for some purposes, despite grouping together functionally diverse strains. We will explain how strong-sense coarse-grainability arises in the model considered here, and show that this property is context-specific: a coarse-graining that works in the organisms’ natural eco-evolutionary context is easily broken if the community is assembled in the non-native environment or if the natural ecological diversity is removed. Finally, we will discuss the extent to which our findings generalize beyond our model.

## I. AN ECO-EVOLUTIONARY FRAMEWORK FOR A HIERARCHICAL DESCRIPTION OF THE INTERACTING PHENOTYPES

In order to study the hierarchy of possible coarsegraining schemes for ecosystems, we need an ecoevolutionary framework that would describe players functionally, by a list of characteristics that can be made longer (more detailed) or shorter (more coarse-grained). In addition, for our purposes we will also want an ability to tune the complexity of the environment, for example, to study the robustness of a coarse-graining between the simplified conditions of a laboratory and the more complex natural environment. In this section, we present our model implementing these two requirements.

### A. The eco-evolutionary dynamics

A given environment presents various opportunities that organisms can exploit to gain a competitive advantage. Imagine a world where all such opportunities or “niches” are enumerated with index *i* ∈ {1 … *L*_∞_}. The notation *L*_∞_ highlights that in general, one expects this to be a very large number, corresponding to a complete (and, in practice, unattainable) microscopic description. A strain *μ* is phenotypically described by enumerating which of these opportunities it exploits, i.e. by a string of numbers of length *L*_∞_ which we will denote *σ*_*μi*_. For simplicity, we will assume *σ*_*μi*_ to be binary (*σ*_*μi*_ ∈ {0, 1}): strain *μ* either can or cannot benefit from opportunity *i*. This will allow us to think of evolution as acting via bit flips 0 →1 and 1 → 0, corresponding to the acquisition or loss of the relevant machinery (“trait *i*”) via horizontal gene transfer events or loss-of-function mutations.

We will assume that the fitness benefit from carrying trait *i* is largest when the opportunity is unexploited, and declines as the competition increases. For a given set of phenotypes present in the community, the ecological dynamics are determined by the feedback between strain abundance and opportunity exploitation (Fig. 1A). Briefly, the strain abundances *N*_*μ*_ determine the total exploitation level *T*_*i*_ ≡ _*μ*_ *N*_*μ*_*σ*_*μi*_ of opportunity *i*. The exploitation level determines the fitness benefit *h*_*i*_ ≡ *h*_*i*_(*T*_*i*_) from carrying the respective trait; we will choose *h*_*i*_(*T*_*i*_) of the form 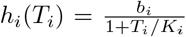. These *h*_*i*_, in turn, determine the growth or decline of the strains. Specifically, we postulate the following ecological dynamics:

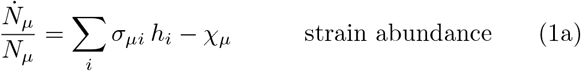

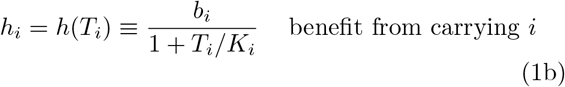

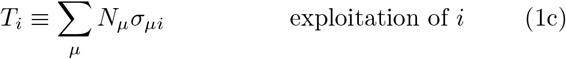

**FIG. 1.**
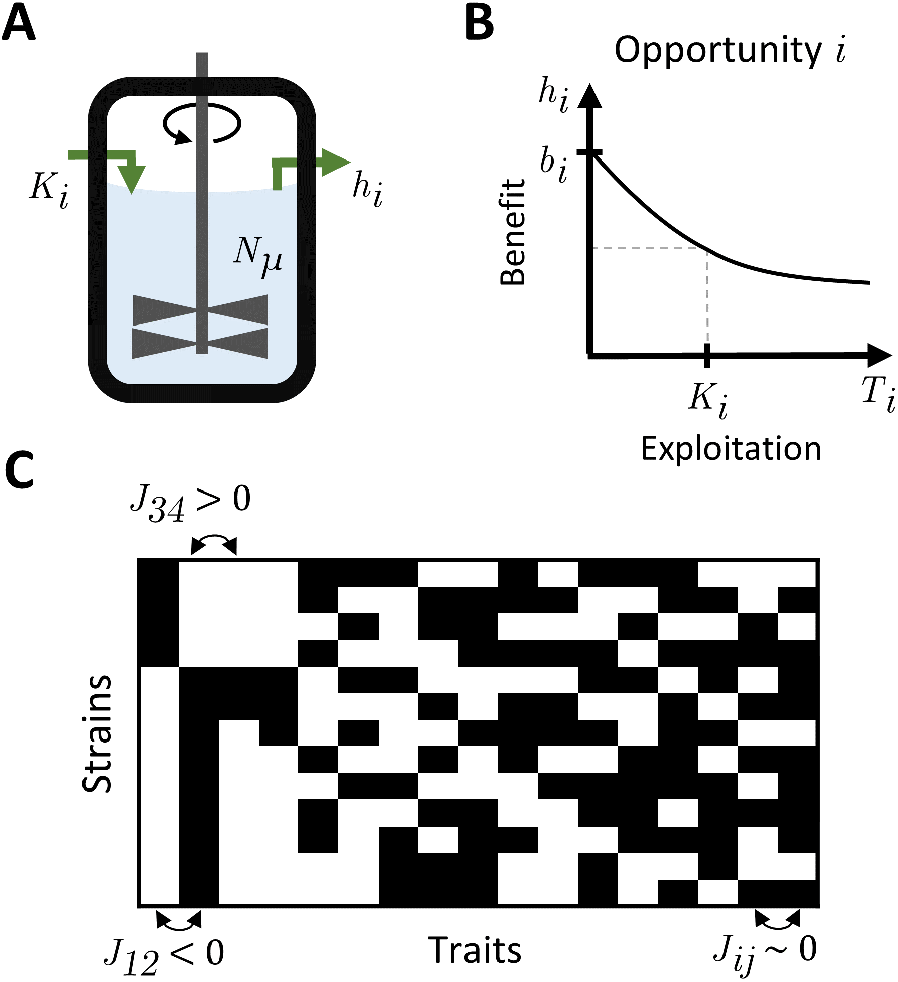
Our eco-evolutionary framework modifies a standard model of resource competition. Organisms engage in ecological competition for limited resources and evolve by gaining or losing traits. Carrying a trait incurs a cost but enables the organism to benefit from the corresponding resource. Here, our novelty is to consider how traits interact with each other. Combinations that interact unfavorably are costly to maintain; as a result, not all phenotypes are competitive. **A:** A metabolic interpretation of our model corresponds to an ecosystem in a chemostat. A set of strains with abundances {*N*_*μ*_} compete for a set of substitutable resources indexed by *i*, e.g., alternative sources of carbon. In this interpretation, *K*_*i*_ correspond to resource supply rates, and *h*_*i*_ are the resource concentrations in the effluent. **B:** For this work, we adopt a more general interpretation where the resources *i* need not be specifically metabolic. Instead, we think of *i* as enumerating any *depletable* environmental opportunities that the phenotypes can exploit, which confer a benefit *h*_*i*_ that declines with exploitation level *T*_*i*_. We parameterize this dependence by the maximum benefit *b*_*i*_ and the carrying capacity *K*_*i*_ (the exploitation level where the benefit is halved); see text. **C:** In our model, phenotypes are binary vectors described by traits they carry. The most competitive phenotypes (rows in the cartoon) are not random, but are shaped by pairwise trait interactions *J*_*ij*_. Strongly synergistic traits (*J*_*ij*_ *>* 0) tend to co-occur, while strongly antagonistic traits (*J*_*ij*_ *<* 0) are likely not carried together. Such structured phenotypes lead to structured ecosystems, as we investigate.

In these equations, the parameters *b*_*i*_ and *K*_*i*_ describe the environment, with *b*_*i*_ being the fitness benefit of being the first to discover the opportunity *i* (at zero exploitation *T*_*i*_ = 0), and the “carrying capacity” *K*_*i*_ describing how quickly the benefit declines as the exploitation level *T*_*i*_ increases (Fig. 1B). The quantities *χ*_*μ*_ are interpreted as the “maintenance cost” of being an organism carrying a given set of traits; more on this below.

The dynamics (1) is basically the MacArthur model of competition for *L*_∞_ substitutable “resources” [28–30]. To these dynamics we add the stochastic arrival of new phenotypes arising through bit flips (“mutations”), as is standard in studies of adaptive dynamics. The combined eco-evolutionary process is simulated using a hybrid discrete-continuous method as described in the SI Appendix A. As presented so far, our eco-evolutionary model is similar to, e.g., Ref. [31]; our key novelty (trait interactions) will be introduced in the next section. We note, however, that typically the interpretation of resources in models like (1) is metabolic [16, 19, 20, 32–35]; for example, *i* might label the different forms of carbon available to a carbon-limited microbial community. Here, we adopt a more general perspective, where *i* labels any depletable environmental opportunity, which need not be specifically metabolic.

As an example, one way for a strain to survive in chemostat conditions is to develop an ability to adhere to the walls of the device [36]. The wall surface is finite, and provides an example of a non-metabolic limited resource. Similarly, being physically bigger, or carrying a rare toxin could be a useful survival strategy, but in both cases the benefit decreases as the trait becomes widespread in the community. Unlike the forms of carbon, which may be numerous but are certainly countable and finite, the list of exploitable opportunities of this kind could be arbitrarily long (*L*_∞_ → ∞), especially when considering the complexity of natural microbial environments. Note that, by construction, our model allows coexistence of a very large number of phenotypes. In many studies, explaining such coexistence is the aim; here, it is our starting point. Rather than asking how a given environment enables coexistence of a diverse community, we start from the observation that natural communities are extremely diverse, interpret this as evidence for the existence of a very large number of (potentially unknown) limiting factors, and ask whether such diversity of types can be usefully coarse-grained.

Modeling fitness benefits as additive [Eq. (1a)] is certainly a simplification. It is also worth noting that the model (1) is special in that it possesses a Lyapunov function [37]; we will return to this point below. Nevertheless, this is a good starting step for our program, namely understanding the circumstances under which coarse-grained descriptions are adequate. Most crucially, a suitable choice of the cost model *χ*_*μ*_ will allow us to naturally obtain communities with an hierarchical structure of trait distributions across organisms mimicking that of natural biodiversity.

### B. A simple cost model leads to hierarchically structured communities

Several studies investigated dynamics like (1) with costs assigned randomly [e.g. 15–20]. Here, we seek to build a model where the phenotypes in the community are not random, but are hierarchically structured, reproducing phenomena such as divergent taxa belonging to identifiable functional groups, the fine-scale strain diversity found within a species, or the notion of “core” and “accessory” traits in a bacterial pangenome [38]. For this, consider the following cost structure:

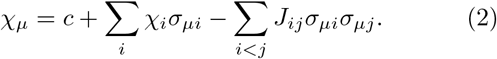

The parameter *c* encodes a baseline cost of essential housekeeping functions (e.g. DNA replication). *χ*_*i*_ is the cost of carrying trait *i* (e.g. synthesizing the relevant machinery); for most of our discussion, we will set *c* = 0.1, and set all *χ*_*i*_ ≡ *χ*_0_ = 0.5 for simplicity. The key object for us is the matrix *J*_*ij*_, which encodes interactions between traits and shapes the pool of viable (low-cost) phenotypes (Fig. 1C). As an example, the enzyme nitrogenase is inactivated by oxygen, so running nitrogen fixation and oxygen respiration in the same cell would require expensive infrastructure for compartmentalizing the two processes from each other; in our model, this would correspond to a strongly negative *J*_*ij*_ (carrying both traits is costly). An example for the opposite case of a beneficial interaction (positive *J*_*ij*_) is a branched catabolic pathway, where sharing enzymes to produce common intermediates reduces the cost relative to running the two branches independently. Crucially, in our model, the parameters *c, χ*_*i*_ and *J*_*ij*_ are the same for all organisms; we will refer to them as encoding the “biochemistry” of our eco-evolutionary world.

We now make our key choice. To set *J*_*ij*_, we generate a random matrix of progressively smaller elements, as illustrated in Fig. 2A. Specifically, we will be drawing the element *J*_*ij*_ out of a Gaussian distribution with zero mean and standard deviation *J*_0_ *f* (max(*i, j*)), with a sigmoidshaped 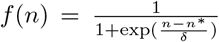 (see Fig. 2B). Throughout this work, we set *J*_0_ = 0.2, *n*^*^ = 10 and *δ* = 3. As we will see, this choice for the interaction matrix *J* implements a hierarchically structured distribution of traits. Intuitively, since high-cost phenotypes are poor competitors, we can think of the interactions *J*_*ij*_ as determining the “sensible” trait associations. For strongly interacting traits only some combinations are competitive, resulting in traits that are mutually exclusive (*J*_*ij*_ *<* 0) or that frequently co-occur (*J*_*ij*_ *>* 0) in low-cost (viable) phenotypes (Fig. 1C). In contrast, a weakly interacting trait can be gained, lost, or remain polymorphic, as dictated by the environment. An example might be a gene encoding a costly pump that enables the organism to live in otherwise inaccessible (toxin-laden) regions of the habitat. Such a trait is “weakly interacting” if the cost of running the pump does not significantly depend on the genetic background. As we will see, our model will naturally give rise to hierarchically structured sets of phenotypes that share some “core” functions but differ in others to form finer-scale diversity, resembling the notions of “core” and “accessory” traits of a bacterial pangenome [38].

**FIG. 2.**
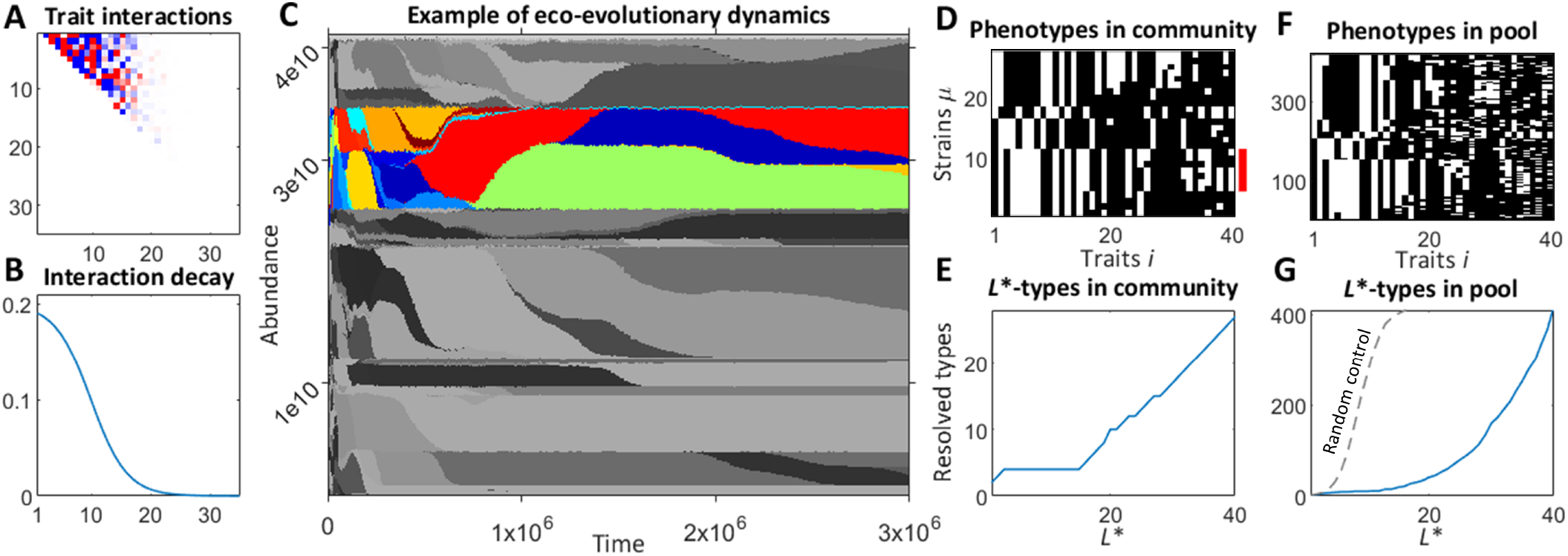
A simple model of trait interactions leads to hierarchically structured ecosystems. **A, B:** In our model, the traits carried by a given phenotype interact with each other to determine its “maintenance cost” (see text). The matrix of pairwise trait interactions *J*_*ij*_ is drawn randomly and is the same for all phenotypes, encoding the “biochemical constraints”; panel A shows an example (*J*_*ij*_ is triangular with one element per trait pair *i* ≠ *j*). We assume an interaction structure such that a few traits interact strongly while others interact weaker and weaker (panel B). **C:** An example of eco-evolutionary dynamics generated in our model. Shading corresponds to different phenotypes. Although new strains continue to emerge and die out throughout the period shown, they can be grouped into several coarse-grained types of approximately stable abundance (one is highlighted in color). **D:** The phenotypes present at the endpoint of the trajectory shown in C. Each of 27 phenotypes is a row of length *L*_*∞*_ = 40 (white pixels are carried traits). The seven highlighted strains are identical in traits 1–24. We will say that they belong to the same “*L*^*\**^-type,” for level of coarse-graining *L*^*\**^ = 24. **E:** The number of *L*^*\**^-types in the community of panel D, shown as a function of *L*^*\**^. At a coarse-grained level, the community appears to consist of only 4 types (one of these is highlighted in C using color); resolving finer substructure requires *L*^*\**^ *>* 15. **F, G:** Same as D, E for a broader set of strains, pooled over *N*_env_ = 50 similar environments. The hierarchical structure is maintained (if the trait matrix were randomized, the number of *L*^*\**^ types would grow exponentially; see the dashed line). Here, we ask: in what sense, if any, could the phenotypic details beyond *L*^*\**^ *≈* 20–25 be coarse-grained away in this model?

### C. Environment defines a strain pool

To build some intuition about the model defined above, consider Figure 2C that shows an example of these eco-evolutionary dynamics for one random biochemistry, and an environment where we set *b*_*i*_ ≡ *b*_0_ = 1 for simplicity, and *K*_*i*_ = *K*_0_ = 10^10^ to set the scale of population size as appropriate for bacteria. The community was initialized with a single (randomly drawn) phenotype. Shading corresponds to distinct phenotypes. The panel illustrates that our framework will allow us to define a form of ecosystem stability where all the original phenotypes may have gone extinct and were replaced by others, and yet at a coarse-grained level the ecosystem structure remains recognizably “the same”. Here, starting from about *t* ≃ 10^5^, the dynamics resemble a stable coexistence of several coarse-grained “species” (one is highlighted in color), whose overall abundance remains roughly stable even as individual strains continue to emerge and die out. To formalize this observation, we need the notion of coarse-grained “*L*^*^-types”, which we will now introduce.

As we continue the simulation, the dynamics converge to an eco-evolutionary equilibrium (a state where the coexisting types are in ecological equilibrium, and no single-bit-flip mutant can invade). In this example, it consists of 27 coexisting phenotypes and is shown in Fig. 2D. Note that, confirming our expectations, it appears to possess a hierarchical structure. The seven highlighted strains are identical over the first 24 components, and differ only in the “tail” (components 25-40). A coarsegrained description that characterizes organisms only by the first *L*^*^ = 24 traits would be unable to distinguish these strains; we will say that these strains belong to the same *L*^*^-type with *L*^*^ = 24. Fig. 2E plots the number of *L*^*^-types resolved at different levels of coarse-graining *L*^*^ (within the community shown in Fig. 2D). For *L*^*^ = 3-15, the number of types remains stable at just 4; the color in Fig. 2C highlights one of them. Beyond *L*^*^ = 15, adding more details begins to resolve additional types, up until *L*^*^ = *L*_∞_ when the number of *L*^*^-types coincides with the total number of microscopic strains.

Of course, when discussing the diversity of strains one expects to find in a given environment, it is important to remember that no real environment is exactly static, and no real community is in evolutionary equilibrium. To take this into account while keeping the model simple, we will consider not a single equilibrium, but a collection of communities assembled in *M*_env_ = 50 similar environments where we randomly perturb the carrying capacity of all opportunities (*K*_*i*_ = *K*_0_(1 + *ϵη*_*i*_), with *ϵ* = 0.1 and *η*_*i*_ are i.i.d. from a standard Gaussian); see SI Appendix A 1. Figure 2F shows the set of strains pooled over the 50 ecosystems assembled in this way. This *strain pool* is the central object we will seek to coarse-grain. We stress that its construction explicitly depends on the environment. (Or, more specifically, the particular random set of *M*_env_ similar environments, but *M*_env_ = 50 is large enough that the results we present are robust to their exact choice.)

As we see in Figure 2F, adding more strains to the pool makes its hierarchical structure even more apparent. Quantitatively, the number of *L*^*^-types (Fig. 2G) grows much slower than if the traits of each phenotype were randomly permuted (the dashed control curve): microscopically, perturbing the environment favors new strains, but at a coarse-grained level, these new strains are variations of the same few types. This is precisely the behavior that we were aiming to capture in our model. Beyond *L*^*^ ≈ 20 − 25, the number of resolved types begins to grow rapidly. Can this diversity be coarse-grained away? Is there a precise sense in which these tail-end traits are “just details”? To answer this question, we must begin by making it quantitative.

## II. COARSE-GRAINING

### A. Methodology for defining coarse-grainability

The *L*_∞_-dimensional description we defined represents the complete list of niches and opportunities present in a natural habitat. Any recreation in the laboratory is simplified, retaining only some of the relevant factors. We will model simplified environments as including resources/opportunities 1 through *L* (Fig. 3A). The parameter *L* represents environment complexity. The other key parameter is the level of coarse-graining detail, *L*^*^ (Fig. 3B). For each *L*^*^, the identity and combined abundance of *L*^*^ types provides a candidate coarse-grained description of the ecosystem. We seek a quantitative metric for assessing its quality.

**FIG. 3.**
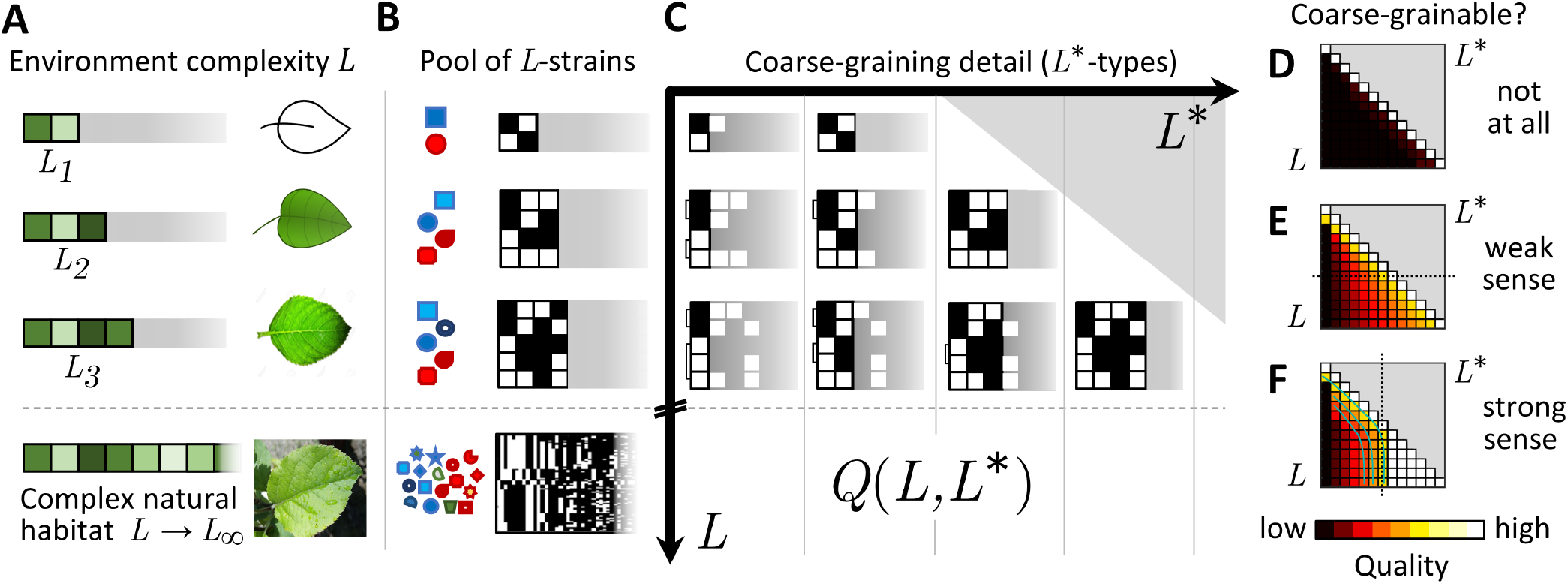
Defining weak and strong coarse-grainability. **A:** The complex natural habitat is modeled as including a large number *L*_*∞*_ of exploitable resources or opportunities. In a laboratory, we can consider a sequence of ever-more-detailed approximations including resources 1, …, *L* (with the remaining ones set to zero). **B:** For each environment, the model describes the pool of strains we expect to encounter (the pool of “*L*-strains”; see Fig. 2F). For a given *L*, the strains are unlikely to carry traits *i* for resources not provided (*i > L*). As environment complexity *L* increases, the pool becomes increasingly diverse. **C:** The set of *L*-strains can be coarse-grained to a varying level of detail *L*^*\**^ *≤ L*. Let *Q*(*L, L*^*\**^) be any quantitative metric (to be defined later) scoring the quality of the *L*^*\**^-coarse-graining in the environment of complexity *L*. At *L*^*\**^ = *L*, the strain diversity is fully resolved (no coarse-graining). The “coarse-grainability” of the ecosystem is encoded in the behavior of *Q*(*L, L*^*\**^) at *L*^*\**^ *< L*. Different metrics *Q* encode different operational definitions of coarse-grainability. **D:** A non-coarse-grainable ecosystem (*sensu* quality metric *Q*). The coarse-graining quality remains poor unless the microscopic strain diversity is fully resolved (at *L*^*\**^ = *L*). **E:** Weak-sense coarse-grainability: in any given environment (a fixed *L*, highlighted), a desired quality can be achieved with a coarser-than-microscopic description (*L*^*\**^ *< L*). **F:** Strong-sense coarse-grainability: the same coarse-graining (a fixed *L*^*\**^, highlighted) provides the desired quality even as the environment complexity is increased.

Ideally, this assessment would be a comparison of performance of two models: one highly detailed, the other coarse-grained, and our test would evaluate the prediction error for a given property of interest. However, what we built is not a coarse-grained *model*, but a hierarchy of coarse-grained variables. These variables could be used to build any number of models, and identifying the most predictive of these is a highly nontrivial task. Inspired by recent experimental work, here we sidestep this problem by proposing an operational approach that evaluates a coarse-graining based on the reproducibility of outcomes for a specified experimental protocol.

We will describe and contrast two protocols, each of which could be seen as verifying the validity of the coarse-graining, and each yielding its own metric of coarse-graining quality *Q*(*L, L*^*^); Fig. 3C. Its “diagonal” (with *L*^*^ = *L*) corresponds to a scenario where the description of strains resolves *all* the traits relevant in a given environment, so must necessarily score as “perfect” under any sensible evaluation criterion. Coarse-grainability is encoded in the behavior of *Q*(*L, L*^*^) with *L*^*^ *< L* (Fig. 3D-F). Consider first the behavior of *Q*(*L, L*^*^) as a function of *L*^*^, with *L* fixed. If we observe that in a given environment, sufficient quality can be achieved already with *L*^*^ *< L*, we will say that the ecosystem is coarse-grainable in the weak sense. For strong-sense coarse-grainability, we ask if the same coarse-grained description continues to perform well even as the environment is made more complex (i.e., instead of fixing *L* and varying *L*^*^, we fix *L*^*^ and vary *L*). Strong-sense coarse-grainability would be a highly desirable property, but *a priori* it is unclear if it is even theoretically possible.

Crucially, these definitions depend on the choice of the operational criterion for assessing coarse-graining validity (the experiment whose results we require to be reproducible). Below, we will show that the same ecosystem can be coarse-grainable in the strong sense under one criterion, and yet not coarse-grainable at all under another.

### B. Operational definitions of coarse-graining quality *Q*(*L, L*^*\**^)

In this section, we describe two “experimental” protocols, each of which could be seen as a sensible test of the quality of a coarse-graining. They will establish two alternative criteria for a coarse-graining to be operationally valid, which we will then contrast.

#### The reconstitution test

One possible criterion is the *reconstitution test*. Drawing a random representative for each of the *L*^*^-types in the strain pool, we seed an identical environment with the representatives we chose, allowing them to reach an ecological equilibrium (Fig. 4B). If the details ignored by the coarse-graining are indeed irrelevant, we would expect such “reconstituted” replicates to all be alike. If the reconstituted communities are found to be highly variable depending on exactly which representative we happened to pick, this will signal that the distinctions we attempted to ignore are, in fact, significant.

**FIG. 4.**
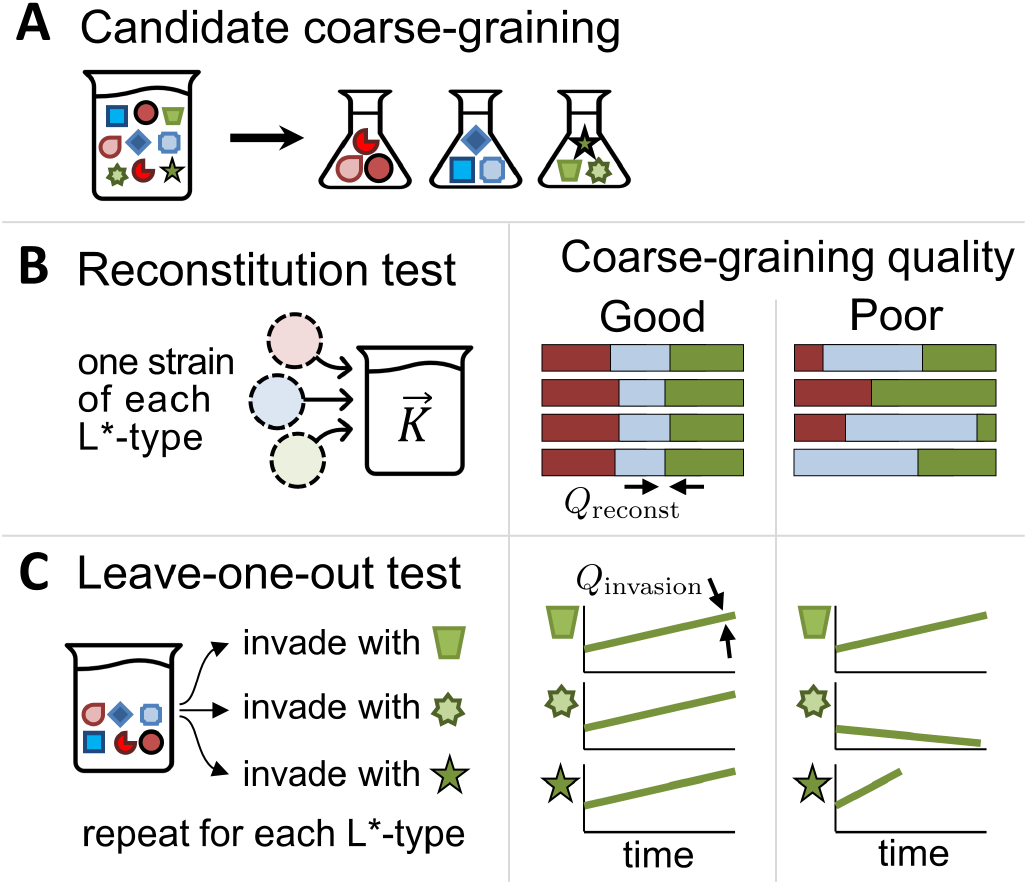
Specific criteria for assessing coarse-graining quality *Q*(*L, L*^*\**^). **A:** In this cartoon, the community is coarse-grained into three operational taxonomic units (OTUs), implemented in our model as *L*^*\**^-types. **B:** The reconstitution test. Under this criterion, grouping strains into coarse-grained OTUs is justified if reconstituting a community from a single representative of each OTU yields similar communities regardless of which representatives we pick. As a quantitative measure, we compare the OTU abundances across replicates. **C:** The “leave-one-out test”. Under this criterion, grouping strains into coarse-grained OTUs is justified if the strains constituting OTU *X* (green in this cartoon) all behave similarly when introduced into a community missing *X*. As a quantitative measure, we compare the invasion rates of the left-out strains.

Quantitatively, for each *L*^*^-type *μ*, let us denote 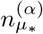 its final relative abundance (i.e., the fraction of total population size) in the reconstituted replicate *α*. The coefficient of variation of 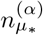 over *α* (denoted 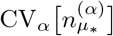) provides a natural measure of variability across replicates. To combine these into a single number, we compute the average such variability over all *L*^*^-types *μ*_*_, weighted by their mean relative abundance across replicates (denoted 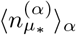):

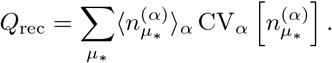

Since the coefficient of variation is, by definition, 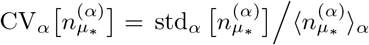, our metric simplifies to 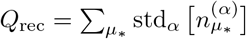. With this definition, a perfect reconstitution would have *Q*_rec_ = 0. Conveniently, this is automatically the case if *L*^*^ = *L* (no coarse-graining).

#### The leave-one-out test

As we will see, the criterion defined above is extremely stringent and is rarely satisfied. In this section, we introduce a weaker version. Instead of the composition of the entire community, we will explicitly focus on one particular property of interest (below, the invasion rate of a strain). Further, instead of requiring the grouped-together strains to be interchangeable in absolute terms, we will ask that they behave similarly *in the context of the assembled community*.

Specifically, for a given scheme grouping strains into coarse-grained types, consider assembling a community missing a particular coarse-grained type *μ*_*_ (the ecological equilibrium reached when combining all the strains in the pool, except those belonging to type *μ*_*_; Fig. 4C). We will judge the coarse-graining as valid if the different strains constituting the missing type *μ*_*_ all behave similarly when introduced into this community. As one example, we can compare their initial growth rates if introduced into the community at low abundance, called henceforth “invasion rate” (other possible choices include the abundance the strain will reach if established, or the level of niche exploitation *h*_*i*_ in the resulting community; these are shown in the SI Fig. S1). If the invasion rates are similar, describing the community as missing the coarse-grained type *μ*_*_ would indeed be consistent. If, however, the invasion rates vary strongly, we will conclude that the features our coarse-graining is neglecting are, in fact, important.

Quantitatively, denote the invasion rate of strain *μ* into a community missing type *μ*_*_ as *r*_*μ,μ\**_. We define

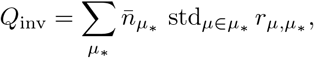

where 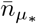 is the relative mean abundance of strains belonging to type *μ*_*_ in the pool, and std_*μ*∈*μ\**_ denotes the standard deviation over all strains belonging to *μ*_*_ weighted by strain abundance in the pool (i.e., a strain’s combined abundance observed across the set of *M*_env_ environments used to define the pool). Once again, at *L*^*^ = *L* we automatically have *Q*_inv_ = 0. Note that this averaging convention (weighted by abundance in the pool) is slightly different from that used in the previous section (using average abundance across the assembled replicates). Using the same convention for both *Q*_inv_ and *Q*_rec_ would not change our results, but would artificially inflate the latter with noise from low-abundance (rare) strains. (For details, see SI Appendix C 2 and Fig. S2.)

To illustrate the difference between the two criteria, consider the statement that a community consisting of *Tetrahymena thermophila* and *Chlamydomonas reinhardtii* cannot be invaded by *Escherichia coli* [39]. What meaning should we ascribe to this statement when phrased in terms of coarse-grained units, rather than specific strains? Under the first criterion, we would require that if we combine any single strain of *T. thermophila*, any strain of *C. reinhardtii*, and any strain of *E. coli*, only the first two would survive. Under the second criterion, we would combine a vial labeled *T. thermophila*, containing the entire diverse ensemble of its strains, with a similarly diverse vial of *C. reinhardtii*, and verify that the resulting community cannot be invaded by any individual strain of *E. coli*.^1^

Note that in our model, the existence of a Lyapunov function [37] means the ecological equilibrium is uniquely determined by the environment and the identity of the competing strains; their initial abundance or the order of their introduction does not matter. While this is a simplification, this property is very useful for our purposes, since any lack of reproducibility between reconstituted communities is then clearly attributable to faulty coarse-graining. In a model where even identical phenotypes could assemble into multiple steady states, distinguishing this variability from the variability due to strain differences would add a layer of complexity to our analysis.

## III. RESULTS

### A. A coarse-graining may be operationally valid despite grouping functionally diverse strains

Throughout this section, we will continue to use an environment with *K*_*i*_ ≡ *K*_0_ and *b*_*i*_ ≡ *b*_0_ (all *L*_∞_ opportunities are equally lucrative). In practice, when approximating a complex environment in the laboratory, we try to capture the most salient features first. Thus, it would have been perfectly natural to instead let *K*_*i*_ and/or *b*_*i*_ decline with *i*; one would expect this to improve coarse-grainability, and this is indeed the case (see SI Appendix S3). The motivation for our choice is two-fold: First, keeping all *K*_*i*_ and *b*_*i*_ the same requires fewer parameters than choosing a particular functional form of decline with *i*. Second, the regime where no niches are obviously negligible will only make it more striking to find that an ecosystem can be not only coarse-grainable, but coarse-grainable in the strong sense.

Fig. 5A plots *Q*_inv_(*L, L*^*^) for the leave-one-out test comparing the invasion rates of different strains falling into the same coarse-grained types. We find that any desired coarse-graining quality can be achieved by a sufficient *L*^*^, and is almost unaffected by *L*. As environment complexity increases and becomes capable of sustaining an ever-growing number of microscopic strains, each *L*^*^ type becomes increasingly diverse. Nevertheless, all the strains in the same *L*^*^-type continue to behave similarly by our invasion-rate-based metric; in other words, under this criterion, the ecosystem is coarse-grainable in the strong sense.

**FIG. 5.**
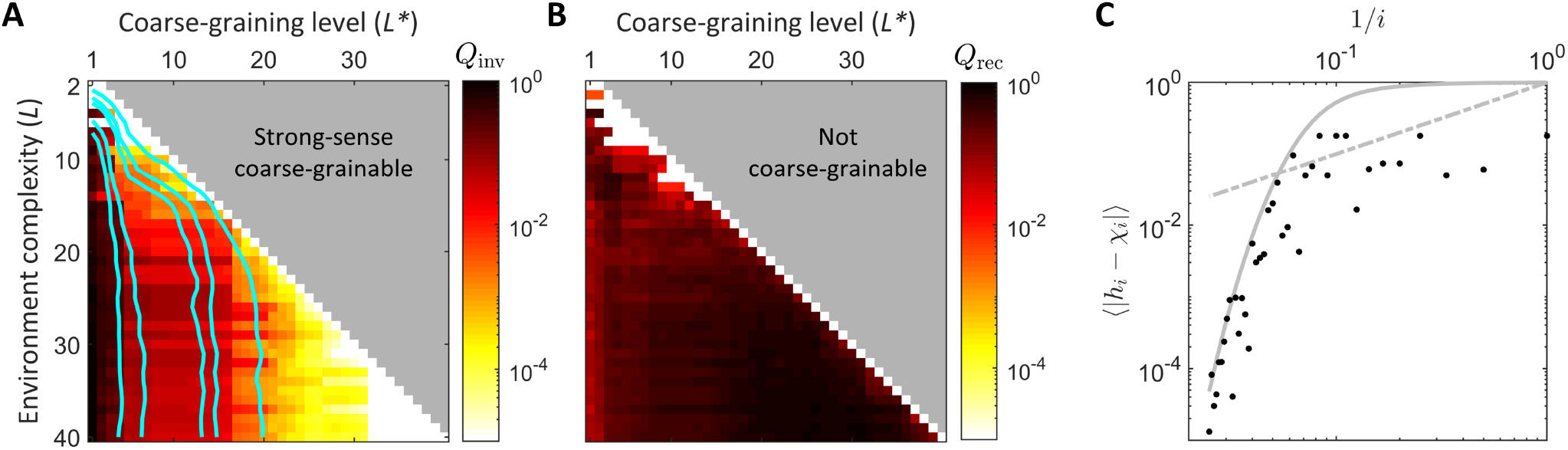
The same ecosystem can be coarse-grainable under one criterion, but not under another. **A:** If coarse-graining quality is evaluated using the leave-one-out test (assessing reproducibility of strain invasion rates), our ecosystem model is coarse-grainable in the strong sense: the acceptable level of coarse-graining, determined by the desired quality score (isolines of *Q*), is robust to environment complexity (compare to Fig. 3F). **B:** In contrast, under the reconstitution test criterion, no amount of coarse-graining is acceptable (compare to Fig. 3D). This comparison shows that a coarse-graining can be operationally valid for a given purpose (panel A) even when the strains it groups together are functionally diverse (panel B). Both heatmaps represent a single random biochemistry, same in both panels. Isolines in A are averaged over 20 biochemistries to demonstrate robustness (see SI Appendix B). **C:** Explaining the origin of strong-sense coarse-grainability in our model. The plot shows the scaling with *i* of |*h*_*i*_ *− χ*_*i*_| (computed for *L*^*\**^ = 30, *L* = 40, and averaged across communities assembled for the leave-one-out test of panel A). The strong-sense coarse-grainability of panel A is ensured whenever the decay is faster than 1*/i* (dashed gray line). Intuitively, this makes the tail-end traits approximately neutral in the assembled community; see text. We expect this scaling to be controlled by the sigmoidal decay of trait interaction magnitude |*J*_*ij*_ |, as confirmed here (solid gray line; same as Fig. 2B but normalized to a maximum of 1 to show the decay of interaction strength rather than their absolute magnitude).

And yet, it would be wrong to conclude that the traits beyond a given *L*^*^ are “negligible” in any absolute sense. This is clearly demonstrated by the reconstitution test (Fig. 5B). If we attempt to reconstruct the community from its members, *every* detail matters: no amount of coarse-graining is acceptable. We will now explain this apparent paradox within our model.

Consider a community at an ecological equilibrium, and let us focus on a particular phenotype *σ* carrying one of the weakly interacting (tail-end) traits *i*_0_: 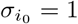. What would be the fitness effect of losing this trait? Losing the benefit 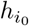 from opportunity *i*_0_ is offset by the reduction in maintenance cost; for a weakly interacting trait, the contribution from the term 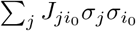 is negligible, and the change in cost is simply 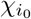. We conclude that the fitness effect of losing the trait is 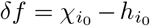. At an evolutionary equilibrium, we would therefore have 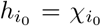 (the “functional attractor” state [20]). When this condition is satisfied, we will say that the opportunity or niche *i*_0_ is “equilibrated”. If a weakly interacting niche is equilibrated, carrying the respective trait becomes approximately neutral.

Here, our community is not at the evolutionary equilibrium; nevertheless, a sufficiently diverse strain pool will similarly ensure that the opportunities corresponding to the weakly-interacting (tail-end) traits become approximately equilibrated: *h*_*i*_ ≈ *χ*_*i*_. For a simpler model where the phenotype costs *χ*_*μ*_ are drawn randomly, the mechanism for this can be understood analytically (the “shielded phase” of Ref. [19]; see also Ref. [40]). Here, the costs are not random, but as long as trait interactions are weak, one expects the behavior to be similar (see SI section S6.2 in Ref. [19]). This expectation is confirmed in simulations. Fig. 5C shows the observed niche disequilibrium *h*_*i*_ − *χ*_*i*_ as a function of 1*/i*. The plot confirms that the tail-end niches (1*/i* → 0) are increasingly well-equilibrated (|*h*_*i*_ − *χ*_*i*_ | decays with *i*). The strong-sense coarse-grainability of Fig. 5A is ensured whenever the decay is faster than 1*/i* (dashed gray line). This is because with this scaling, the sum of contributions from the omitted tail-end traits is bounded (see SI Appendix E). The analytical argument of Ref. [19] leads us to expect the disequilibrium to be controlled by the decaying typical magnitude of interactions |*J*_*ij*_| (solid gray line). If the tail-end niches are equilibrated, carrying the respective traits becomes approximately neutral, and the ability of a strain to invade is entirely determined by its phenotypic profile over non-equilibrated niches, explaining the observations of Fig. 5A. We conclude that in our model, the strong-sense coarse-grainability is a consequence of the faster-than-1*/i* decay of interaction strength in Fig. 3B.

Crucially, however, this approximate neutrality applies only in the environment created by the assembled community, and does not mean that the distinctions are functionally negligible. For instance, consider the (Lotka-Volterra-style) interaction term for a given pair of strains *μ≠ν*:

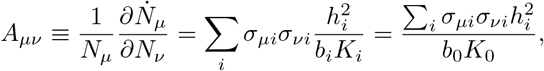

where we substituted *b*_*i*_ ≡ *b*_0_ and *K*_*i*_ ≡ *K*_0_ for our environment. Even when tail-end niches are equilibrated with *h*_*i*_ ≈ *χ*_*i*_ = *χ*_0_, we find that each of them contributes equally to the interaction term: no detail is negligible.

This argument directly relates the observed effect to the distinction between a trait that is truly neutral, and one that is effectively neutral in the assembled community only. A truly neutral trait, one incurring almost no cost and bringing almost no benefit, would have *h*_*i*_ → 0 and its contribution to the interaction term *A*_*μν*_ would indeed be small. And indeed, if we repeat our analysis for a scenario where both *b*_*i*_ and *χ*_*i*_ decline with *i*, we find that neglecting the tail-end traits becomes an adequate coarse-graining also for the reconstitution test (see SI, Fig. S3).

The conclusion from contrasting Fig. 5A and 5B is worth emphasizing. In the example we constructed, the coarse-grained description is valid *sensu* panel 5A. This means that, for instance, we can meaningfully say that “a community assembled of OTU#1 and OTU#2 can be invaded by OTU#4”. We can even measure, e.g., the invasion rate, and be assured that it is quantitatively reproducible, with a bounded error bar, across the many strains that constitute OTU#4 at the microscopic level. Despite all this, the *interaction* between the OTUs as coarse-grained units is not actually definable: any specific pair of strains of OTU#1 and OTU#4 may interact differently with each other, as is indeed observed experimentally [11].

Our focus on reproducibility of *L*^*^-type abundances across replicates is inspired by the experiments Ref. [14]. To complete this parallel, we should mention that besides inoculating the same environment with a set of similar inocula, as we did for our reconstitution test (*cf* Fig. 4B), one could also use the same inoculum to seed a set of similar environments. To implement this in our model, we use the strain pool constructed as described in section I C to inoculate a set of environments with slight variations in the carrying capacities *K*_*i*_ ≈ *K*_0_ drawn from a Gaussian distribution of width *ϵ* = 0.1. This is meant to represent the unavoidable variability present in any experimental replicates of the “same” environment *K*_*i*_ ≈ *K*_0_, which can affect fitness even when subtle [41]. After assembling the replicate communities, we find that community composition is more reproducible at coarser levels of description (Fig. 6B,C), consistent with the experimental observations of Goldford et al. [14], and with the interpretation of this pattern as resulting from functional redundancy within coarse-grained types [6, 42].

**FIG. 6.**
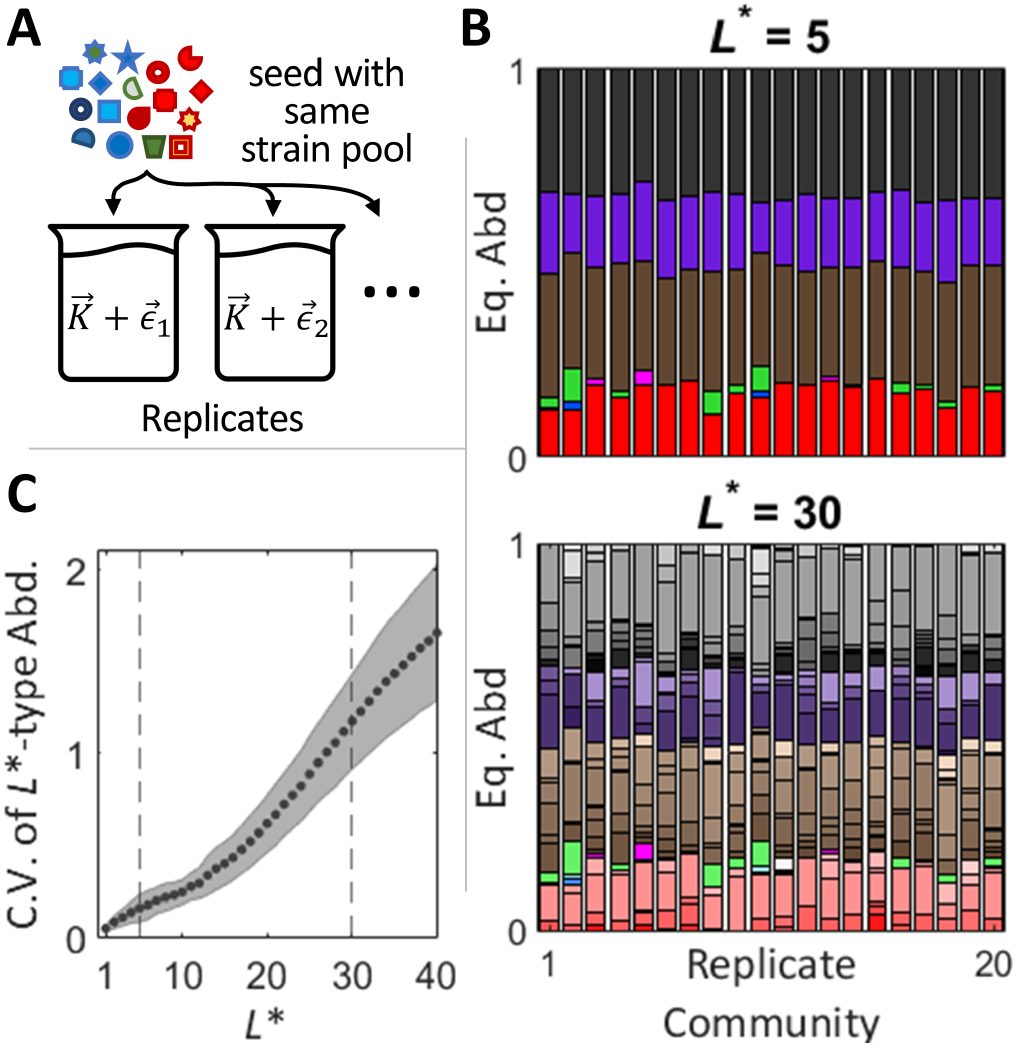
Replicate communities assembled in similar environments are more reproducible at coarser level of description. **A:** A set of similar environments 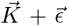 (each carrying capacity modified by 10% Gaussian noise) is inoculated with the same strain pool, and brought to eco-logical equilibrium. **B:** Equilibrium relative abundances of coarse-grained *L*^*\**^-types across 20 replicates, shown for two levels of coarse-graining. A coarser description (*L*^*\**^ = 5; 7 resolved types) is more reproducible, consistent with experimental observations [14]. **C:** The variability of coarse-grained descriptions increases with level of detail. Variability is measured as the average coefficient of variation (C.V.) in relative abundance of an *L*^*\**^-type over 100 replicates, weighted by *L*^*\**^-type mean relative abundance across replicates. Dashed lines mark *L*^*\**^ = 5, 30 shown in B. Datapoints and shading show mean *±* SD over 20 random choices of biochemistry {*J*_*ij*_}. All simulations performed with *L* = 40.

### B. Using non-native strain pool reduces coarse-grainability

The previous section describes a mechanism by which strain diversity can aid coarse-grainability. As we explained, in our model ecosystem the diverse set of strains contained within the coarse-grained units was able to successfully equilibrate the weakly interacting niches, rendering them effectively neutral and leading to the behavior shown in Fig. 5A. However, for this to occur, the strain pool diversity needs to be derived from a sufficiently similar set of environments, as we will now show.

To see this, we repeat the leave-one-out analysis of Fig. 5A, except now we inoculate the same test environment of complexity *L* = 40 (using *K*_*i*_ = *K*_0_, *b*_*i*_ = *b*_0_ as before) with strain pools derived from *other* environments that are increasingly dissimilar to it. Specifically, following the procedure described in section I C, we generate strain pools in environments with *K*_*i*_ = *K*_0_(1+*ϵη*_*i*_), where *η*_*i*_ are drawn from the standard normal distribution, and *ϵ* is the parameter we vary. (The *b*_*i*_ are left at *b*_*i*_ = *b*_0_ for simplicity.) The results are presented in Figure 7, which shows the performance of different *L*^*^-coarse-grainings under the leave-one-out test.

**FIG. 7.**
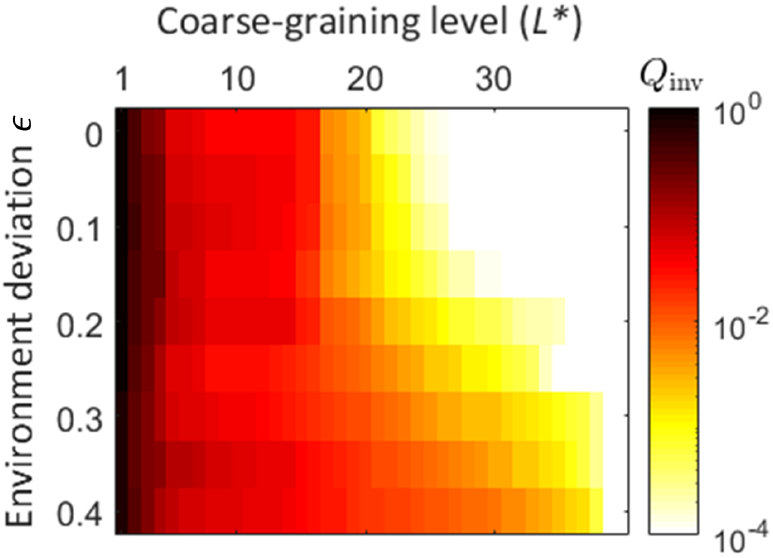
A coarse-graining scheme works best when the environment is populated by the native strain pool. The same test environment as in Fig. 5A is inoculated with strain pools that evolved in environments increasingly further away (see text). The coarse-graining quality is assessed by leave-one-out experiments, and shown as a function of *L*^*\**^ and environment deviation *ϵ* from the test condition. *L* is fixed at *L* = 40 for comparison with the last row of Fig. 5A. As the environments for generating strain pools are modified, the traits that were previously negligible can no longer be coarse-grained. The same random biochemistry as in Fig. 5A was used, and each pixel is averaged over 20 random environments.

At *ϵ* = 0, this is identical to the protocol of Fig. 5A. We see that describing phenotypes by 20 traits is sufficient for the invasion rates of grouped-together strains to be consistent within an error bar of *Q <* 10^−2^. However, as *ϵ* is increased, and the strain pools we use are derived from increasingly distant environments, the same coarse-graining becomes insufficient. Instead, a substantially higher level of coarse-graining detail *L*^*^ is required to maintain the desired quality. In summary, we find that in our model, a coarse-graining scheme works best when the environment is populated by the native strain pool.

## IV. DISCUSSION

The interface of statistical physics and theoretical ecology has a long and highly influential tradition of studying large, random ecosystems, starting from the work of May [43]. The key insight of this approach is that patterns that are typical to some *ensemble* of ecosystems are more likely to be generalizable and reproducible than the details specific to any one realization. However, the choice of the ensemble (and in particular, adding constraints relevant for natural ecosystems) can affect predictions significantly [44–48]. Which predictions of random-interaction models are robust to introducing more realistic structures, and conversely, which phenomena cannot be explained without invoking structural constraints, is an active area of research [49].

Resource competition models—one of the simplest frameworks explicitly linking composition to function— offer a highly promising context to begin addressing these questions, with much recent progress. For example, it was recently shown that cross-feeding interactions structured by shared “rules of metabolism” (but otherwise random) can reproduce a surprising range of experimental observations [14, 16, 17]. This work made it possible to begin disentangling which experimental observations can be seen as evidence for nontrivial underlying mechanisms, and which can be reproduced already in the simplest models.

In this work, we presented a simple framework that allows generating random ecosystems with community structure as a tunable control parameter. Instead of postulating a fixed architecture, such as a number of discrete “families” of phenotypes [17], we use a biologically motivated approach to derive it from functional tradeoffs, parameterized by a matrix of trait-trait interactions *J*. Simple (few-parameter) choices for *J* generate communities with complex structures, including hierarchical architectures which, at least superficially, appear to mimic those of natural biodiversity. Perhaps the most immediate benefit from such a framework would be to help develop new ways to quantify the highly multi-dimensional concept of “community structure” across scales, such as, for example, the structure of microbial pangenomes.

In this spirit, here we used this framework to quantify the notion of coarse-grainability. We proposed a way to operationally define the quality of a coarse-grained description based on the reproducibility of outcomes of a specified experiment. We demonstrated that an ecosystem can be coarse-grainable under one criterion, while also not at all coarse-grainable under another.

Specifically, one way to approach the coarse-graining problem is to only group together the individuals that are to a sufficient extent interchangeable. This is the criterion we introduced as a “reconstitution test”, and is the criterion implicitly assumed by virtually all compositional models of ecosystem dynamics [50]. However, experimental evidence [3–5, 8, 11] suggests that unless we are willing to resolve types differing by as few as 100 bases, this criterion is likely violated in most practical circumstances. It is certainly violated when grouping strains into taxonomic species or families [6, 13, 51–53]. One expects, therefore, that explaining the practical successes of such descriptions would require a different definition of what makes a coarse-graining scheme adequate.

We proposed that this can be achieved with only a subtle change to the criterion: namely, by requiring that the grouped strains be approximately interchangeable not in all conditions, but in the conditions created by the assembled community itself. As long as the strains we study remain in a diverse ecological context, and as long as this diversity is derived from a sufficiently similar environment, we find that the coarse-grained description can be consistent in the sense that the strains grouped together possess similar properties of interest (e.g., invasion rate, post-invasion abundance).

In this paper, we focused on a case where the traits were differentiated only by the strength of their interactions, which established a unique hierarchy among them (a clear order in which to include them in the hierarchy of coarse-grained descriptions). In the more general case, the trait cost *χ*_*i*_, or the trait usefulness in a given environment (*b*_*i*_, *K*_*i*_) will set up alternative, potentially conflicting hierarchies. We expect the model to have a rich phenomenology in this regime, which we have not considered here. Another obvious limitation of our analysis is that our model includes only competitive interactions. A simple way to extend our framework would be to include cross-feeding interactions; we leave this extension for future work.

Our analysis introduced a distinction between weak-sense and strong-sense coarse-grainability based on whether the performance of a coarse-graining scheme is robust to increasing the environment complexity. We explained how strong-sense coarse-grainability arises in our model, linking it to a previously described phenomenon, namely that a sufficiently diverse community may “pin” resource concentrations (here, the exploitation of environmental opportunities) at values that are robust to compositional details [17, 19, 32, 40]. Tracing its origin makes it clear that strong-sense coarse-grainability in our model is only as good as the assumption that the cost of carrying weakly-interacting traits is independent of the phenotypic background. Whether this assumption is ever a good approximation in natural ecosystems remains to be seen. Still, our argument provides an explicit mechanism for how coarse-grainability can not only coexist, but may in fact be facilitated by diversity.

The fact that strong-sense coarse-grainability is at least theoretically possible is intriguing also for the following reason. Throughout this work, we interpreted *L* as indexing a sequence of ever-more-complex environments (e.g., a minimal medium with 1 carbon source; a complex mixture of carbon sources; resuspended leaf mush; an actual leaf). An alternative perspective, however, is to think of a single environment of interest and take *L* to be the level of detail at which it is modeled. Any model we could ever consider, however detailed, is necessarily incomplete. Consider the example of the human gut: how important is the exact geometry of the gut epithelium? the effect of peristalsis and flow on small-scale bacterial aggregates? the exact role of the vast diversity of uncharacterized secondary metabolites [54, 55]? It seems plausible that the complete list of factors shaping this ecosystem includes many we will never even know about, let alone include in our models. Our analysis raises an intriguing—though at this point, purely speculative—question of whether the tremendous diversity of natural ecosystems might afford our models some unexpected degree of robustness to such unknown details.

In conclusion, there are many reasons to believe that analyzing a species in artificial laboratory environments might be of limited utility for understanding its function or interactions in the natural environment [56]. Usually, however, the concern is that the laboratory conditions are too simple, and in reality, many more details may matter. Here, we use our model to propose that, at least in some conditions, the opposite can be true: understanding the interaction of two strains in the foreign conditions of the Petri dish may require a much more detailed knowledge of microscopic idiosyncracies. Removing individual strains of a species from their natural eco-evolutionary context may eliminate the very reasons that make a species-level characterization an adequate coarse-graining of the natural diversity.

All simulations were performed in MATLAB (Mathworks, Inc.). The associated code, data, and scripts to reproduce all figures in this work are available at Mendeley Data [57].

## ACKNOWLEDGMENTS

We thank J. Grilli, C. Holmes, R.S. McGee, and C. Strandkvist for helpful discussions. This research was supported in part by NSF Grant No. PHY-1748958, the Gordon and Betty Moore Foundation Grant No. 2919.02, and the Kavli Foundation.

## Appendix A: Simulating eco-evolutionary dynamics

The eco-evolutionary world in which the dynamics take place is described by the environment (constant in time), the biochemistry (also constant in time), and the state of the ecosystem (dynamically evolving). At any given moment of time, the state of the ecosystem is described by the following information: (1) The identity of each of the phenotypes, described microscopically as vectors of length *L*_∞_; (2) The current abundance (population size) of each of these phenotypes. All simulations are performed in the *L*_∞_-dimensional world of complete microscopic detail; the environments of reduced complexity *L* are implemented by zeroing out the environmental niches from *L* + 1 onward.

At the level of individual bacteria, for any moment in time, the next “event” to occur will be one of the following: (1) an individual dies; (2) an individual divides, giving rise to an identical sibling; or (3) an individual divides, giving rise to a mutant sibling. Of course, an individual-based simulation is both impractical and unnecessary; instead, we think of these dynamics as a combination of purely ecological updates of phenotype abundances (which can be modeled with continuous ODEs), and discrete dynamics whereby some strains go extinct, and others are introduced into the population by mutation.

To implement such discrete events, the standard way is to employ a Gillespie scheme [58]. A slight complication here is that when overlaid with ecological dynamics, the rates of such Gillespie events become time-dependent (a mutation favorable right now may cease to be so as ecological dynamics continue). However, this complication is easily resolved using standard methods for implementing a hybrid stochastic-deterministic Gillespie scheme [59]. Briefly, instead of drawing the “time to next event”, one must draw a probability threshold, and propagate the continuous dynamics while integrating the rate of an event to occur, up until that accumulated probability crosses the threshold (see [60] for an introduction that is both short and intuitive). As a result, to describe our simulation we just need to define how the state-dependent rates of such events are computed.

To do so, we adapt for our purposes the results of Ref. [61]. From the evolutionary standpoint, the candidate new strain is a mutant that has a chance of escaping drift and become *established* in the population. The probability of becoming established is proportional to the mutation rate, the population size of the parent strain, and the fitness effect *δf* of the mutation (i.e. the growth rate^2^ of the candidate new strain). Once a strain is established, the stochastic effects become negligible and its subsequent dynamics can be modeled deterministically.

The simulation can be summarized with the following pseudocode:

1. For all single mutants of existing strains, determine the rate at which they would establish in the population, as explicit functions of the abundances of extant strains.
2. Propagate ecological dynamics “for an appropriate length of time” as per standard technique [59].
3. Pick the lucky new strain among the beneficial first mutants; add it to the community at a population size 1*/δf* ; the population of the parent strain is, for consistency, reduced by the same amount.
4. Remove any strain whose abundance is below some predetermined threshold and is declining. (In our simulations, this threshold is set at relative abundance 10^−6^.)
5. Repeat until the required simulation time has elapsed.

The mutation rate we use is *μ* = 10^−10^ per individual per unit time. For context, recall that population size is set by the carrying capacity of environmental niches *N* = 10^10^ (see Fig. 2C), so our choice corresponds to *μN* = 1. However, as explained below, the mutation rate parameter plays only a minor role in our analysis.

### 1. Evolutionary equilibrium is not required

In our analysis of coarse-graining schemes, the community we study is never technically at an evolutionary equilibrium. We explicitly constructed our procedure to avoid making such an unrealistic postulate. Specifically, note that we assemble our strain pool from evolutionary equilibria obtained in *similar* environments, but we then study the interaction of these strains in the original, unperturbed environment, where the condition of evolutionary equilibrium was never imposed.

Of course, we then observe that a sufficiently diverse set of strains derived from sufficiently similar environments assembles into a community that is very close to the evolutionary equilibrium also in the original environment, and this proximity is largely responsible for the behaviors reported in this work. This, however, is not a caveat, but a feature, as we expect the same to be largely true for real communities as well: the large diversity of strains is quite plausibly sufficient to populate the available niches without requiring *de novo* mutations, relying exclusively on the standing variation.

### 2. Mutation rate is not a key parameter

A corollary of the previous point is that for specifically our purposes here, mutation rate is not a key parameter of our model. Indeed, we only invoke evolution when constructing the strain pool, but each of the combined states is an evolutionary equilibrium, which in this model is guaranteed to be unique. One small caveat is that our evolutionary process is simulated at a finite resolution, and considers first-mutants only. As a result, the trajectory could get stuck in a *locally* non-invadeable equilibrium rather than the unique true one, something that would be enhanced by setting the mutation rate too low. In this way, the evolutionary stochasticity (and thus the mutatiton rate) does technically play a weak role, but we found it to be essentially irrelevant for the parameters used here (specifically, running replicate eco-evolutionary trajectories from random initial phenotypes generated virtually indistinguishable final states).

One artifact that occasionally arises when constructing a strain pool in an environment of complexity *L* occurs when some obtained phenotypes are identical in all traits *i* ≤ *L* but are distinguishable in traits past *L*, which correspond to those resources/opportunities that have been set to zero available benefit (*h*_*i*_ = 0 by *b*_*i*_ being set to zero for *i > L*; see Appendix A above). Note that in our model, it is rare but perfectly reasonable for phenotypes to carry traits that offer no environmental benefit; this occurs whenever the contribution of trait interactions *J*_*ij*_ provides a net reduction of maintenance cost. (In other words, surviving strains need not be identically 0 past *i* = *L*.) However, what should be true is that an *L*^*^ type with *L* = *L*^*^ can only have one representative: in any environment of complexity *L <*≤ *L*^*^, the lowest-cost member in any *L*^*^-type is strictly superior to all others in its *L*^*^-type, and would outcompete them. In practice, the inferior strains are occasionally retained in some replicates due to the evolutionary process only considering first mutants. This has no effect on eco-evolutionary dynamics beyond a slight change to the abundance of the respective type; however, if left uncorrected, it would lead to artifacts in evaluating coarse-graining quality due to the artificially inflated diversity within an *L*^*^-type. To correct for this, we add the following step when generating the strain pool: after collecting together all phenotypes from the ensemble of similar environments, we check for any *L*^*^ types that contain more than one strain at *L* = *L*^*^ and, if so, remove this artifact by retaining only the superior (lowest-cost) phenotype.

## Appendix B: Averaging in figures

All heatmaps shown in figures correspond to a single random biochemistry. The small amount of “graininess” seen in the heatmaps is caused by the quirks of exactly when the *L*^*^-coarse-graining is able to resolve new subtypes. It is shown as is, with no additional averaging or smoothing. In contrast, the isolines overlaid on plots are meant to capture the qualitative trends transcending the quirks of a given biochemistry. They are computed by repeating the same analysis for 20 random biochemistry realizations: the 20 heatmaps are averaged, smoothed with a 1-pixel-wide Gaussian kernel, and the isolines are picked as contour lines of the result.

The only exception to this procedure is Fig. 7 in the main text which investigates the effect of perturbing the environment. In this figure, each pixel is an average over 20 random perturbations (of magnitude specified by epsilon) for a single biochemistry.

## Appendix C: Metrics for Coarse-graining Quality

### 1. Examples of other ecological properties and their compatibility with *L*^*\**^-coarse-graining

As described in the main text, the leave-one-out scheme judges a grouping of strains into coarse-grained OTUs as justified if the strains constituting OTU *X* all behave similarly when introduced into a community missing *X*. However, the “similarity of behavior” could itself be assessed by a variety of criteria. In the main text, we focused on comparing the invasion rates of the left-out strains. Some equally interesting alternatives include, for instance, the abundance reached by the invading strain (which might be zero if the strain cannot invade), or the niche occupancy in the resulting community. In each case, we perform the same weighting procedure as used in the main text when defining the leave-one-out test (*Q*_inv_).

#### Invading strain abundance

In this example, we denote *n*_*μ,μ\**_ the relative final abundance reached by strain *μ* after being introduced in a community missing the *L*^*^-type *μ*_*_ (by construction, *μ* is a representative of *μ*_*_). Similar to the reconstitution test (for abundances), we look at the variability of *n*_*μ,μ\**_ over all *μ* ∈ *μ*_*_, and define the coarse-graining quality *Q*_abd_ by the weighted average of this quantity over all *L*^*^-types:

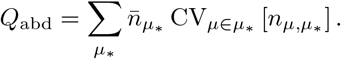

#### Niche occupancy post-invasion

Here, instead of looking at a compositional property (abundance of a particular coarse-grained type), we ask how similar the effect of the invading strains is on the post-invasion *function* of the community as a whole. The functional properties are encoded in community-wide niche occupancy *T*_*i*_ — or, equivalently, in the niche benefit *h*_*i*_ at equilibrium (the benefit from carrying trait *i*). Since the environment is kept fixed, *T*_*i*_ and *h*_*i*_ are directly related. The reason we choose *h* over *T* is because in a metabolic interpretation of the resources (community in a chemostat), *h*_*i*_ is directly measurable as resource concentration in the effluent.

Quantitatively, let 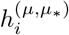 be the *h* vector of the equilibrium community after strain *μ* invades the community missing the *L*^*^-type *μ*_*_ (again, *μ* is a representative of *μ*_*_). One might expect computing the average component-wise standard deviation of this vector (across all *μ* ∈ *μ*_*_) to be an appropriate measure of variability, but doing so artificially attenuates any variation by including “equilibrated” niches with *h*_*i*_ ≈ *χ*_*i*_ (see main text and Appendix E) and thus have vanishing variation. We therefore focus only on the variability in the first component 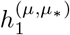. Similar to the leave-one-out invasion rate analysis, we then compute the weighted average of this quantity over all *L*^*^-types:

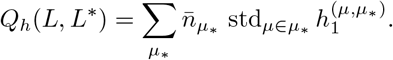

These two new measures of coarse-graining quality, combined with the two considered in the main text, give four metrics *Q*(*L, L*^*^) encoding four different questions of interest. We chose them to span both compositional and functional properties of a community. How amenable are these four questions to coarse-graining?

The answer is presented in Fig. S1. The panels are ordered by “degree of coarse-grainability” as defined in Fig. 3D-F. Our findings are consistent with the recurring observation in the experimental literature that functional properties tend to be more reproducible than compositional ones [6, 14, 51]: the hierarchy of the four questions presented in figure S1 can be summarized as saying that ecosystems are coarse-grainable in the strong sense when a coarse-graining is evaluated based on functional properties.

Indeed, the invasion rate of a missing strain (panel A) is basically a functional property of the pre-invasion community (the growth rate of a strain is determined by the environmental conditions constructed by the rest of the community). Next comes the niche occupancy (panel B) — a functional property assessed post-invasion. Next is the abundance reached by the invading strain (panel C), a compositional property that we expect to be distinctly less coarse-grainable (indeed, recall how strain-strain interactions are strongly affected by tail-end niches). However, the reconstitution test is still last in the list (panel D), epitomizing the goal of a bottom-up compositional description and requiring the knowledge of *all* microscopic details.

### 2. Weighing by Pool Abundance versus Replicate Community Abundance

Recall that the reconstitution scheme (Fig. 4B) evaluates a grouping of strains into coarse-grained OTUs based on the ability of the coarse-graining to precisely reconstitute replicate communities from representatives of each OTU. If the replicate communities are similar in composition, then we deem the coarse-graining to be an adequate grouping. Specifically, we quantify this in terms of a weighted average coefficient of variation in type relative abundance (*n*_*μ\**_) at ecological equilibrium across replicates (*Q*_rec_). In the main text, we choose to perform the weighting of each type by its mean relative abundance over replicates, rather than by its mean abundance in the pool (as used in the leave-one-out scheme). This choice avoids artificially increasing *Q*_rec_ with noise from rare, low-abundance types (Fig. S2). To see this mathematically, consider a low-abundance *L*^*^-type, 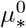 that is rarely observed across *M* replicate communities: say, 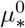 has abundance *ϵ* in only 1 replicate. When *M* is large, the coefficient of variation is approximately 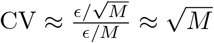.

**FIG. S1.**
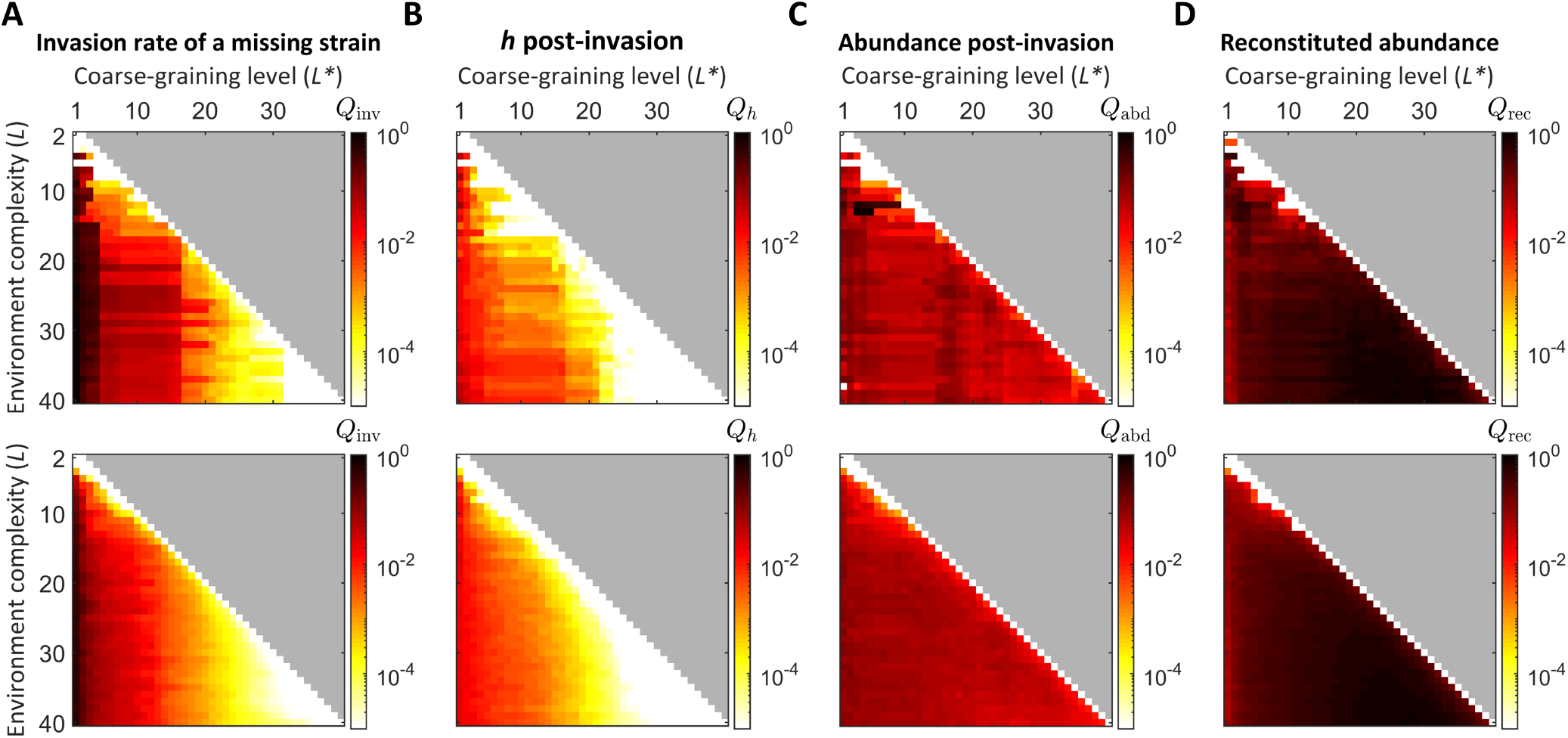
Four questions whose compatibility with the *L*^*\**^ coarse-graining scheme ranges from excellent to non-existent. Top row of heatmaps show coarse-graining qualities for a single random biochemistry, while those in the bottom row show averages over 20 random biochemistries from which the isolines shown in Fig. 5 are computed. **A:** Invasion rate of a missing strain; coarse-grained description sufficient. **B:** Niche occupancy post-invasion; coarse-grained description sufficient. **C:** Abundance reached by a strain post-invasion; coarse-graining compatibility is poor. **D:** Reconstitution test (requires functional equivalence of strains); no coarse-graining is possible. Note that the coarse-graining quality metrics shown in panels A-C are all assessed in the framework of the leave-one-out test as defined in the text.

In the case of weighing by replicate-mean abundance, *Q*_rec_ is simply the sum of standard deviations (as noted in the main text) to which the rare type contributes 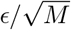, which is very small if *ϵ* is small compared to *M*. In the alternative case of weighing by pool-mean abundance, the contribution from the rare type’s C.V., 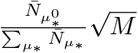, may or may not be small depending on the relative abundance of the strains that constitute type 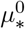 in the pool, inflating the noise from its rare appearances.

**FIG. S2.**
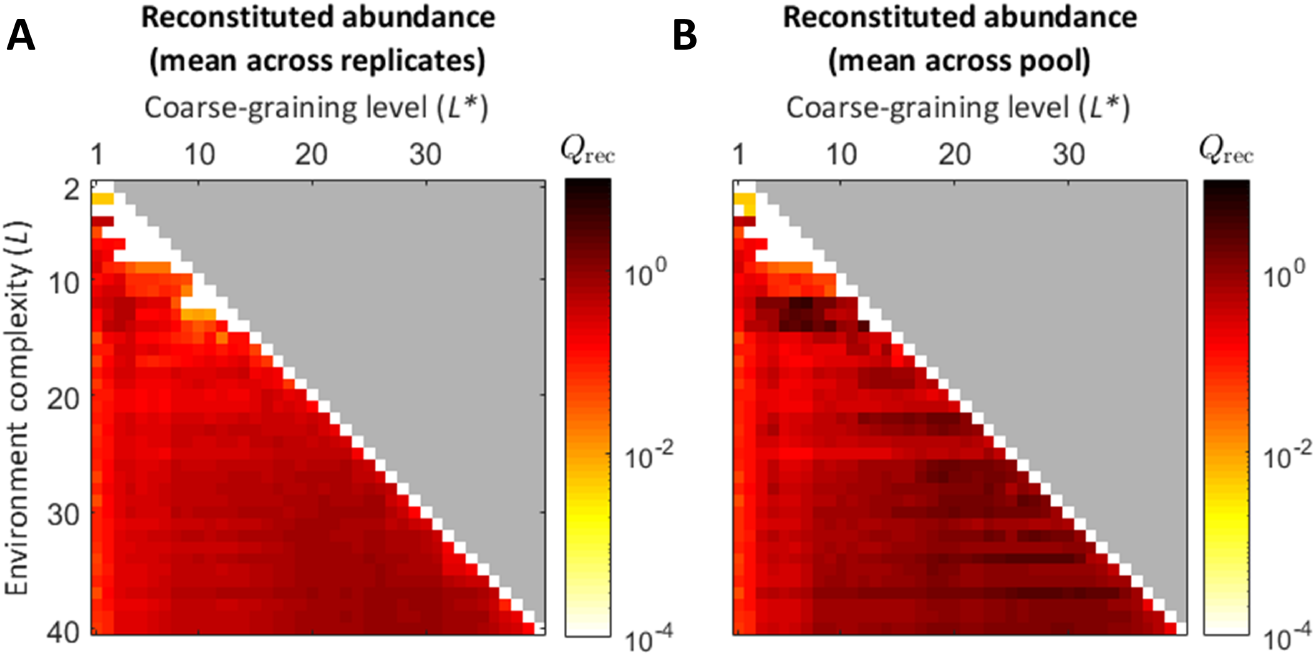
Comparison of weighting *L*^*\**^-type coefficient of variations (C.V.) in *Q*_rec_ by a type’s mean abundance either as observed across replicate reconstituted communities (**A**) or in constructing the strain pool (**B**). The latter weighting inflates the noise of rare (low abundant) strains causing an increase in *Q*_rec_, which indicates larger typical variability within *L*^*\**^-types. Each panel shows the same biochemistry seed as used in Fig. 5.

## Appendix D: Coarse-graining truly neutral traits

One simple sanity-check of our framework is to make the tail-end niches *i* → *L*_∞_ not just weakly interacting, but genuinely close to neutral. A distinction that makes almost no difference in either cost or benefit should surely be negligible for all questions, including the “reconstitution test”. To test this, we repeat the analysis for a model where we use the same declining sigmoid-shaped function *f* (*n*) to scale down not only the trait interactions (cf. Fig 2A, B), but also the trait cost *χ*_*i*_ and benefit *b*_*i*_. The result is shown in Fig. S3. As expected, we find that all four example properties and criteria we consider are now coarse-grainable in either the strong or weak sense (this figure is to be compared with Fig. S1).

**FIG. S3.**
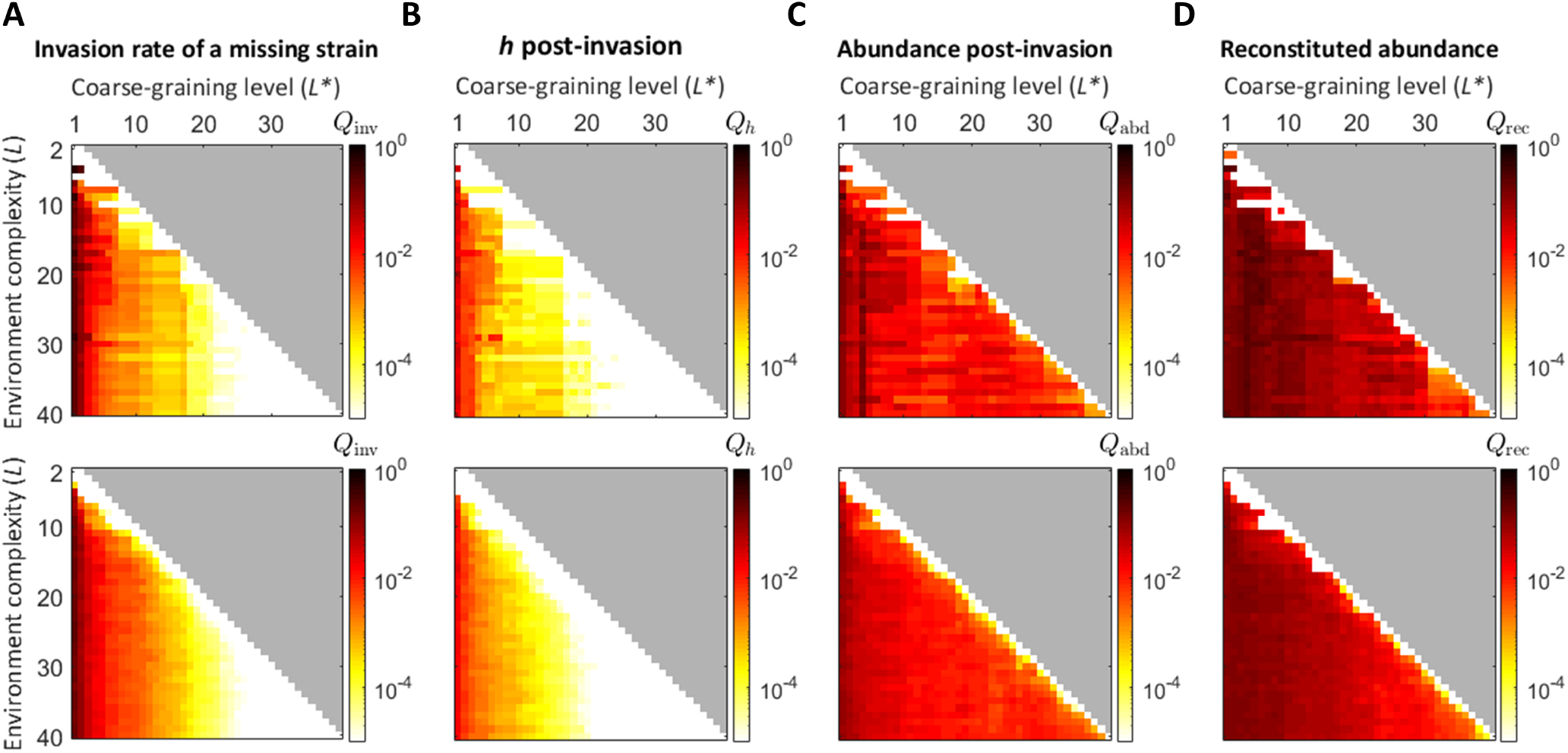
Reproducing Fig. S1 for a model where tail-end niches are not just weakly interacting, but are also increasingly neutral (bring almost no benefit *b*_*i*_ → 0 and incur almost no cost *χ*_*i*_ → 0); see text. As expected, phenotypic diversity in the corresponding traits can be adequately coarse-grained away no matter the criteria (compare to Fig. S1). Again, top row of heatmaps show coarse-graining qualities for a single random biochemistry, while those in the bottom row show averages over 20 random biochemistries.

## Appendix E: Effectively neutral traits and asymptotic scaling of *Q*_inv_(*L, L*^*\**^)

As described in the main text, with respect to the invasion rates of strains in leave-one-out experiments, our ecosystem is coarse-grainable in the strong sense, meaning that a given *L*^*^-coarse-graining maintains a desired quality even as more diversity is resolved with increasing environment complexity *L*. In other words, the variability of invasion rates between strains within *L*^*^-types can be made arbitrarily small with a sufficient level of coarse-graining (*L*^*^), independent of the environment complexity. This appendix presents the formal analysis underlying this result.

Recall that *Q*_inv_(*L, L*^*^) measures the typical invasion rate variability between strains within *L*^*^-types so that *Q*_inv_ = 0 indicates a perfect coarse-graining, which trivially occurs when *L*^*^ = *L*. Put mathematically, for invasion rate to be a strong-sense coarse-grainable property, it must satisfy the following criterion: for some sufficient *L*^*^ *< L*, there exists some finite (possibly *L*^*^-dependent) constant *M* (*L*^*^) such that *Q*_inv_(*L, L*^*^) *< M* (*L*^*^) for any *L*. We first analyze only the invasion rate variability between strains within some *L*^*^-type, from which the same analysis applies to any *L*^*^-type.

The invasion rate of strain *μ* is determined by the pre-invasion niche availabilities *h*_*i*_ set by the community missing type *μ*_*_:

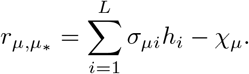

For a given *L*^*^, all strains within a type *μ*_*_ possess identical traits up to *i* = *L*^*^ so that the differences in invasion rates follow from differences in traits *i > L*^*^. Let then *r*_.,*μ\**_ denote the identical contributions to invasion rate from traits *i* ≤ *L*^*^ in order to isolate the variable part:

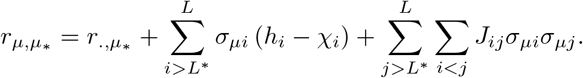

For the coarse-grainability criterion to hold, we need to show that each of these sum s converge as *L* → ∞. Dealing first with the trait-trait interaction piece (*J*_*ij*_ terms), we define the finite sum *S*_*j*_ ≡ Σ_*i<j*_ *J*_*ij*_*σ*_*μi*_ and take *L* → ∞ so that the double sum is converted into a series of finite sums,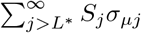. In order for this series to converge, we must have |*S*_*j*_| fall off faster than 1*/j*. To determine the scaling of the *S*_*j*_, note that the terms of each finite sum are Gaussian distributed with mean ⟨*J*_*ij*_⟩ = 0 and sigmoidally decaying variance 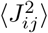 (see main text Sec. I B). Therefore, each finite sum is essentially a random walk of terms that on average sum to 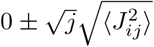, where for large *j* (as *L* tends to infinity) the scatter goes like 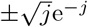 due to the sigmoidal decay of trait interaction strength. With the *S*_*j*_ thus falling off exponentially, we indeed have convergence:

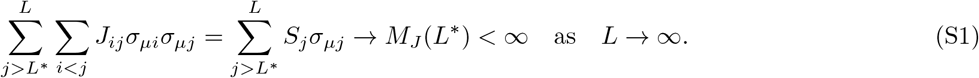

Similarly, for the non-interaction piece to converge as *L* → ∞, we must have |*h*_*i*_ − *χ*_*i*_| ∼ 1*/i*^*α*^ with *α >* 1. Fig. S4 shows the scaling of the typical cost-benefit deviation ⟨|*h*_*i*_ − *χ*_*i*_|⟩ with 1*/i*, where the average is over (left-out) types with the same weighting as used in the main text. Plotted with the simulation data are visual guides that show the scaling of the convergence condition (1*/i*, dashed 1:1 line) and the scaling of the trait interaction sigmoid used to generate *J*_*ij*_ (solid gray line). We see that for a sufficient level of coarse-graining (*L*^*^ = 30, black points) the data indeed falls off faster than 1*/i*, verifying that

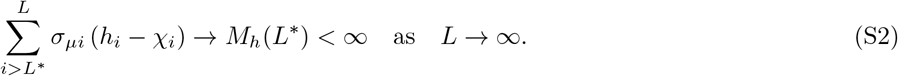

This result follows from two contributing factors: both (1) weak *J*_*ij*_ and (2) phenotypic diversity in the tail-end traits. Ref. [19] has demonstrated that when a community is seeded with a sufficiently diverse strain pool consisting of unstructured, purely random phenotypes (*σ*_*μi*_) the community exhibits a collective phase where strain abundances will adjust to drive fitness benefits to match costs, *h*_*i*_ ≈ *χ*_*i*_ (effectively neutral traits in the so called “S-phase”). To see the presence of this phase in the leave-one-out context, first note that the pool of phenotypes generated within our eco-evo framework are essentially random and unstructured in those traits that are weakly interacting (small *J*_*ij*_) as observed in the example shown in Fig. 2F. Second, as *L*^*^ increases and more strains are resolved, fewer strains are removed in leaving out an *L*^*^-type, increasing the diversity of the remaining community. Fig. S4 demonstrates the effect of this point: for small *L*^*^, the community to be invaded is not diverse enough for weakly interacting traits to be considered effectively neutral (*h*_*i*_ *≉ χ*_*i*_) and therefore cannot be ignored (*L*^*^ = 3, yellow points), unlike the large *L*^*^ regime (*L*^*^ = 30, black points).

Together, (S1) and (S2) imply that the typical invasion rate variation between strains is asymptotically bounded by some constant *M* (*L*^*^) that is independent of *L*, enabling coarse-grainability in the strong sense.

**FIG. S4.**
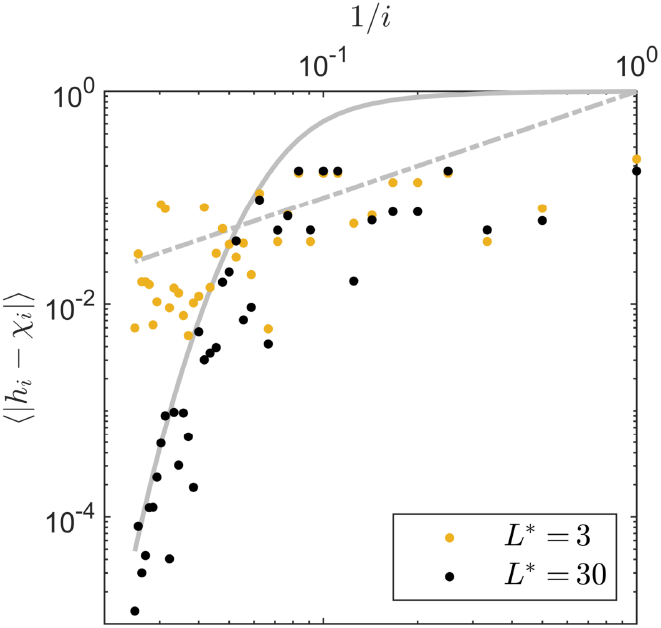
Asymptotic scaling of the typical cost-benefit deviation (|*h*_*i*_ *− χ*_*i*_|) in the leave-one-out scheme. For weakly interacting traits (large *i*), the corresponding niche benefits become effectively neutral when sufficient diversity is present in the community missing an *L*^*\**^-type (black points; *L*^*\**^ = 30), vanishing exponentially (cf. trait interaction decay |*J*_*ij*_| plotted as a solid gray line) such that their sum converges as described in expression S2. When *L*^*\**^ is too small (e.g., *L*^*\**^ = 3; yellow points), insufficient diversity remains after removing an *L*^*\**^-type such that the sum will typically no longer converge (terms scaling like dashed 1*/i* line) and coarse-graining quality is poor. Both coarse-grainings are for an environment of complexity *L* = 40 and the same biochemistry as in Fig. 5.

## Footnotes

1 Although we use this as an example here, we should note that in the original reference [39] this statement was not actually meant as a species-level claim, but indeed referred to the three specific strains used in the experiment.

2 Note that in many evolutionary models, the fitness effect is computed relative to the (ever-increasing) mean fitness in the population. For us here, fitness is not an abstract property that could increase arbitrarily, but is directly defined as a growth rate; and limited resources automatically ensure that the population-mean growth rate is zero.

